# Neutrophils initiate the destruction of the olfactory epithelium during SARS-CoV-2 infection in hamsters

**DOI:** 10.1101/2022.03.15.484439

**Authors:** Bourgon Clara, St Albin Audrey, Ando-Grard Ophélie, Da Costa Bruno, Domain Roxane, Korkmaz Brice, Klonjkowski Bernard, Le Poder Sophie, Meunier Nicolas

## Abstract

The loss of smell related to SARS-CoV-2 infection is one of the most prevalent symptoms of COVID-19. It is now clear that this symptom is related to the massive infection by SARS-CoV-2 of the olfactory epithelium leading to its desquamation. However, the molecular mechanism behind the destabilization of the olfactory epithelium is less clear. Using golden Syrian hamster, we show here that while apoptosis remains at a low level in damaged infected epithelium, the latter is invaded by innate immunity cells. By depleting the neutrophil population or blocking the activity of neutrophil elastase-like proteinases, we reduced the damage induced by the SARS-CoV-2 infection. Surprisingly, the impairment of neutrophil activity led to a decrease of SARS-CoV-2 infection levels in the nasal cavity. Our results indicate a counterproductive role of neutrophils leading to the release of infected cells in the lumen of the nasal cavity and thereby enhanced spreading of the virus.

## Introduction

Loss of smell (anosmia) is a major symptom of COVID-19 pandemic. With omicron’s increased transmission, hundreds of thousands of people per day still get infected worldwide. Despite omicron’s reduced anosmia prevalence (UK Health Security Agency Technical Briefing Jan 14, 2022), loss of smell will likely affect millions more (Vaira et al., 2021; von Bartheld et al., 2020) and 10% of anosmic patients might not recover their sense of smell 6 months after the disease onset (*3*, *4*). The full olfactory recovery could even take up to one year and some patients may never recover their sense of smell (*5*). A recent study estimates that in the US about 720,000 people actually suffer from chronic olfactory disorder related to COVID-19 (*6*). The loss of smell negatively impacts life quality by disrupting feeding behavior potentially leading to malnutrition; and by exposing to food poisoning and to inhalation of dangerous chemicals (*7*). In severe and persistent cases, anosmic patients could possibly suffer from chronic depression (*8*). It is thus crucial to understand the cellular basis of anosmia.

Olfaction starts in the olfactory epithelium (OE) which contains olfactory sensory neurons (OSNs) surrounded by supporting cells called sustentacular cells (SCs). Both cell types are regenerated regularly due to multipotent basal cells (*9*). Among these cells, only SCs express significantly ACE2 and TRPMSS2 required for SARS-CoV-2 cellular entrance (*10*–*12*). In a previous study, we observed in the golden Syrian hamster model that SARS-CoV-2 infects massively the sustentacular cells in the OE leading to its desquamation and olfactory neurons loss (Bryche et al., 2020). Although very rarely OSNs may be infected by SARS-CoV-2 (*15*), a recent study in humans concludes that anosmia arises primarily from infection of sustentacular cells of the OE followed by the disruption of OE integrity without OSN infection (Khan et al., 2021b).

Previous studies in rodents have concluded that more than 90% of the OE needs to be destroyed to impair olfactory-mediated food-finding behaviour (*16*). This suggests that a large area of the OE has to become dysfunctional to experience anosmia in COVID-19. How does SARS-CoV-2 achieve such an unprecedented success in dismantling the sense of smell, compared with previous coronavirus and influenza pandemics? Several studies reported that most cells of the infected OE including OSNs undergo apoptosis (*17*–*20*) leading to desquamation. A similar phenomenon has been reported during influenza infection (*21*) and has been considered for SARS-CoV-2 as a defence mechanism to limit a potential invasion of the central nervous system by pathogens using the olfactory route (*22*). Alternatively, innate immune cells could trigger directly the desquamation of the OE through inflammation as observed in the lung (*23*). Indeed, innate immune cells invade massively the SARS-CoV-2 infected OE (*13*). Iba1 is a marker of microglia/macrophages (*24*) which are the most studied innate immunity cells in the nasal cavity (*21*). Iba1^+^ microglial cells ensure viral clearance by phagocytizing viral particles and infected cells (*25*, *26*) and can induce cell death as observed in the hippocampus using the Theiler’s virus model of encephalitis (*27*). As the OE is not protected by the blood brain barrier, neutrophils and monocytes/macrophages classically recruited during the early event of inflammation could also be involved in the OE damage following SARS-CoV-2 infection (*28*, *29*). Neutrophils are well known for their ability to induce tissue damage, notably through the release of elastase-like proteinases (*30*, *31*) as well as the production of reactive oxygen species (ROS) by the myeloperoxidase (MPO) and formation of toxic neutrophil extracellular traps (*32*, *33*). Macrophages are known for their ability to phagocyte pathogens, produce cytokines and activate other immune cells (*34*). Although they are involved in the regeneration of the OE (*35*, *36*), they can also lead to tissue damage during viral infections notably through NLRP3 inflammasome activation and metalloproteinases activity (*29*). In this study, we focused on the early events following SARS-CoV-2 infection of the nasal cavity to explore the mechanism of the unusually extensive OE damage following SARS-CoV-2 infection.

## Material and methods

### Study design

The study was performed to understand the cellular mechanisms leading to the SARS-CoV-2 induced damage in the OE using hamsters as an animal model. Hamsters experiments were planned in accordance with the principles of the 3Rs (replacement, reduction, and refinement). Body weight and animal behaviour was monitored before and during the experiments. Different parameters in the nasal cavity were measured by quantitative polymerase chain reaction (qPCR) and by immunohistochemistry. Leucocyte numeration was performed directly by an automated analyser. SARS-CoV-2 replication was measured *in vitro* to evaluate a potential inhibition by the drugs used to modulate neutrophils activity. Sample size for each experiment is indicated in figure legends. During analysis, all data points were included except because of technical failure to process the sample. Animals were randomized to the experimental groups. All analyses were performed blindly of the treatment.

### SARS-CoV-2 isolates

Experiments were carried out with SARS-CoV-2 strain BetaCoV/France/IDF/200107/2020, which was isolated by Dr. Paccoud from the La Pitié-Salpétrière Hospital in France. This strain was kindly provided by the Urgent Response to Biological Threats (CIBU) hosted by Institut Pasteur (Paris, France), headed by Dr. Jean-Claude Manuguerra. Cell culture experiments were performed with the SARS-CoV-2 strain France/IDF0372/2020 kindly provided by Sylvie van der Werf.

### Animals

Fifty-six 8 weeks-old male hamsters were purchased from Janvier’s breeding Center (Le Genest, St Isle, France). Animal experiments were carried out in the animal biosafety level 3 facility of the UMR Virologie (ENVA, Maisons-Alfort); approved by the ANSES/EnvA/UPEC Ethics Committee (CE2A16) and authorized by the French ministry of Research under the number APAFIS#25384-2020041515287655. Infection was achieved by nasal instillation (40 μL in each nostril with 5.10^3^ TCID50 of SARS-CoV2 strain BetaCoV/France/IDF/200107/2020) on anesthetised animals under isofluorane. Seven mock-infected animals received only Dulbecco’s minimal essential medium.

For neutrophil depletion experiments, hamsters were injected intraperitoneally with either PBS or 150 mg/kg and 100 mg/kg of cyclophosphamide (CAS: 6055-19-2; PHR1404; Sigma Aldrich) at 3 and 1 days before SARS-CoV-2 infection. Half of the animals were sacrificed at 1 dpi (days post infection) and the other half at 2 dpi (n=4 in each group).

To inhibit neutrophil elastase-like proteases, we used a synthetic cathepsin C inhibitor (IcatC^XPZ-01^; (*37*) diluted in 10 % (2-Hydroxypropyl)-β-cyclodextrin (CAS: 128446-35-5; C0926; Sigma-Aldrich) suspended in citrate buffer 50 mM at pH=5 (vehicle) as described previously (*38*). Hamsters were injected intraperitoneally twice a day with either vehicle (n=4) or IcatC_XPz-01_ at 4.5 mg/kg for 10 days before infection by SARS-CoV-2 (n=4). Animals were sacrificed at 1 dpi.

For all experiments except IcatC_XPZ-01_ treatment, the head was divided sagitally into two halves, of which one was used for immunohistochemistry experiments. Nasal turbinates were extracted from the other half for qPCR analysis. Only histological analysis was performed on tissues from IcatC_XPZ-01_ treatment experiments.

### Immunohistochemistry and quantifications

The immunohistochemistry analysis of the olfactory mucosa tissue sections was performed as described previously in mice (*39*). Briefly, the animal hemi-heads were fixed for 3 days at room temperature in 4% paraformaldehyde (PFA) and decalcified in Osteosoft (Osteosoft; 101728; Merck Millipore; Saint-Quentin Fallavier; France) for 3 weeks. Blocks were cryo-protected in 30% sucrose. Cryo-sectioning (12 μm) was performed in coronal sections of the nasal cavity, perpendicular to the hard palate in order to examine the vomeronasal organ (VNO), olfactory epithelium (OE), Steno’s gland and olfactory bulb. Sections were stored at −80 °C until use.

Non-specific staining was blocked by incubation with 2% bovine serum albumin (BSA) and 0.05% Tween. The sections were then incubated overnight with primary antibodies directed against SARS Nucleocapsid protein (1/500; mouse monoclonal; clone 1C7C7; Sigma-Aldrich), ionised calcium-binding adapter molecule 1 (Iba1) (1/500; rabbit monoclonal; clone EPR16588; Abcam), myeloperoxidase protein (MPO) (1/500; rabbit monoclonal; clone EPR20257; Abcam), CD68 (1/200; rabbit polyclonal; PA1518; Boster), cleaved caspase 3 (C3C) (1/200; rabbit polyclonal; #9661; Cell signalling), G_olf_ (1/300; rabbit polyclonal; C-18; Santa Cruz) and Olfactory Marker Protein (OMP) (1/500; goat polyclonal; 544-10001, Wako). Fluorescence staining was performed using goat anti-mouse-A555; goat anti-rabbit-A488 and donkey anti-goat-A546 (1/800; Molecular Probes A21422; A11056; A11008 respectively; Invitrogen; Cergy-Pontoise; France).

Images were taken at x100 magnification using a 1X71 Olympus microscope equipped with an Orca ER Hamamatsu cooled CCD camera (Hamamatsu Photonics France; Massy; France). Whole section images were taken at x50 magnification using a Leica MZ10F Fluorescent binocular microscope.

To assess olfactory epithelium damage, we used a global score from 1 to 9 based on the integrity of the OE. To evaluate the correlation of apoptosis and innate immune cell presence with damage, OE areas were divided in two groups: undamaged areas (damage score equal to 1 or 2) and damaged areas (damage score between 5 and 9). Apoptosis level, infiltration of immune cells in the OE and its underlying lamina propria were quantified as the percentage of the area positive for C3C (cleaved caspase 3); Iba1 (macrophages/microglia), CD68 (activated bone-marrow-derived macrophages) and MPO (neutrophils). For each animal, the percentage of stained OE was averaged over 4 distinct areas in the beginning of olfactory turbinates at 1 dpi and in the medial part of the nasal cavity containing Steno’s gland and NALT at 2 dpi.

In cyclophosphamide and in IcatC_XPZ-01_ experiments, we examined two independent sections of nasal turbinates (separated by 500 μm) in the middle of the nasal cavity containing NALT and Steno’s gland. For each section, we set a global score from 1 to 9 for neutrophil infiltration based on the overall presence of MPO signal in nasal mucosa and in nasal cavity lumen. For each section, we also measured the total infected area of the OE, the area of desquamated cells in the lumen (based on Hoechst nuclear staining) and the percentage of infected desquamated cells in the lumen (based on N protein immunostaining).

All quantifications were made with ImageJ (Rasband, W.S., ImageJ, U.S. National Institutes of Health, Bethesda, Maryland, USA, http://imagej.nih.gov/ii/, 1997–2012) to threshold specific staining.

### RNA extraction and RT-qPCR analysis

Total RNA was extracted from frozen nasal turbinates using the Trizol-chloroforme method as described previously (*39*). Oligo-dT first strand cDNA synthesis was performed from 5 μg total RNA with iScript Advance cDNA Synthesis Kit for RT-qPCR (Bio-Rad; #1725038) following manufacturer’s recommendations. qPCR was carried out using 125 ng of cDNA added to a 15 μL reaction mix. This reaction mix contained 10 μL iTaq Universal Sybr Green SuperMix (BioRad; #1725124), and primers at 500 nM (sequences in **Supp. Table 1)**. The reaction was performed with a thermocycler (Mastercycler ep Realplex, Eppendorf). Fluorescence during qPCR reaction was monitored and measured by Realplex Eppendorf software. A dissociation curve was plotted at the end of the fourty amplification cycles of the qPCR to check the ability of theses primers to amplify a single and specific PCR product.

Quantification of initial specific RNA concentration was done using the ΔΔ*Ct* method. Standard controls of specificity and efficiency of the qPCR were performed. The mRNA expression of each gene was normalized with the expression level of G3PDH. A correction factor was applied to each primer pair according to their efficiency (*40*).

### Measure of antiviral activity of cyclophosphamide and cathepsin C inhibitor against SARS-CoV-2 in cell culture

Vero E6 cells (CRL-1586, ATCC maintained at 37 °C; 5% CO_2_) were seeded at 2.10^4^ cells per well in a 96-well plate in Dulbecco’s Modified Eagle’s Medium, 5% foetal bovine serum (FBS-12A, Capricorn Scientific, Clinisciences). For cyclophosphamide antiviral activity evaluation, cells were treated with 0.15 mg/mL (corresponding to the maximum dose potentially present in hamsters) or 0.45 mg/mL cyclophosphamide diluted in sterile PBS. For Icat_CXPZ-01_ antiviral activity evaluation, cells were treated with 4.5 μg/mL (corresponding to the maximum dose potentially present in hamsters) or 13.5 μg/mL IcatC_XPZ-01_ diluted in 10% dextrin, citrate buffer 50 mM, pH=5. Cells were treated with vehicle as control (n=6 for each condition). All treatments were performed one hour prior to infection with SARS-CoV-2 strain France/IDF0372/2020 at 5.10^3^ pfu per well diluted in DMEM, 10% Fetal Bovine Serum. Loss of cell viability reflecting the efficiency of viral infection was measured 3 days after infection by adding 100 μL Cell Titer-Glo reagent to each well (CellTiterGlo Luminescent Cell Viability Assay, Promega #G7571), according to the manufacturer’s protocol. Cell luminescence of each well was then quantified using an Infinite M200Pro TECAN.

### Statistical analysis

All comparisons were made using Prism 5.0 (GraphPad). Statistical significance between groups was assessed using non-parametric Mann Whitney tests. For correlation analyses, we used Spearman non-parametric test. Error bars indicate the SEM. Detailed information on statistical test used, sample size and P value are provided in the figure legends.

## Results

### Apoptosis occurs after cell desquamation following SARS-CoV-2 infection of the olfactory epithelium

We previously observed that as soon as two days following nasal instillation of SARS-CoV-2 in Syrian gold hamsters, the sustentacular cells of the olfactory epithelium were massively infected along with strong cellular loss and cellular debris filling the lumen of the nasal cavity (*13*). In order to understand the events leading to this desquamation, we chose to focus on the early stages of infection at 1 and 2 dpi. To evaluate the importance of apoptosis in the damage of the olfactory epithelium following SARS-CoV-2 infection, we measured the level of cleaved caspase 3 signal in uninfected animals, and in infected zones of the OE that were either intact or damaged (**Fig. 1**). Basal level of apoptosis occurring in the OE was not increased in either zone at 1 or 2 dpi (**Fig. 1D**). However, we observed a strong cleaved caspase 3 signal co-localizing partly with desquamated cell in the lumen of the nasal cavity. The cleaved caspase 3 signal in the lumen of the nasal cavity was increased 5- and 14-fold compared to the OE at 1 and 2 dpi respectively, which was statistically significant at 2 dpi (Mann Whitney, *p*=0.0286) and nearly significant at 1 dpi (*p*=0.0525).

**Figure 1:**
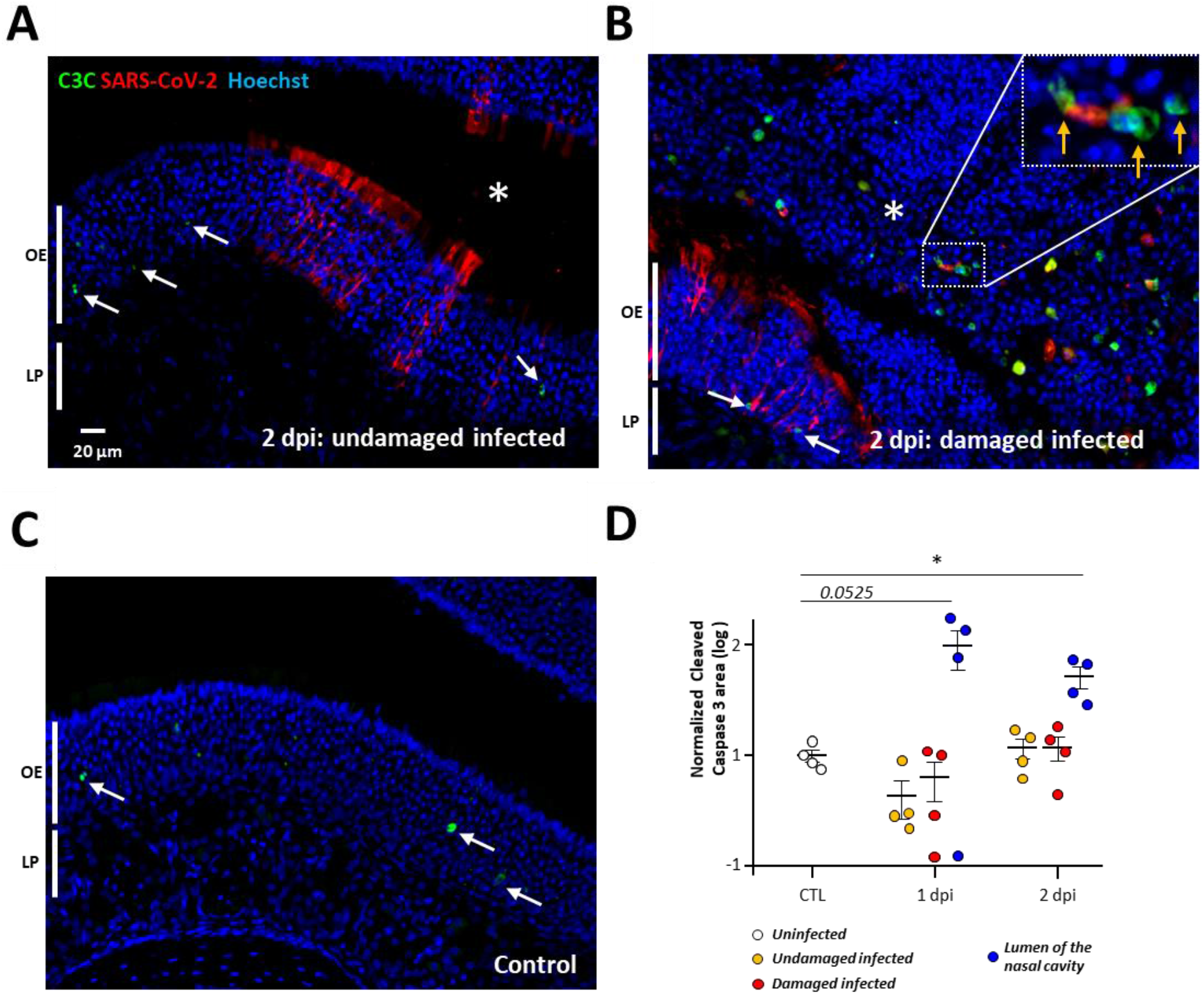
Apoptosis occurs in desquamated cells in the lumen of the nasal cavity following SARS-CoV-2 infection but not in the olfactory epithelium. Representative images of an infected intact (**A**), infected damaged (**B**) and non (**C**) infected area of the olfactory epithelium at 2 days post infection (dpi). Apoptotic cells in the olfactory epithelium are indicated by a white arrow (OE; Olfactory Epithelium / LP Lamina Propria). The lumen of the nasal cavity is indicated by a white asterisk and is filled with cells some of which colocalize in their nucleus cleaved caspase 3 signal (orange arrow). (**D**) Cleaved caspase 3 signal in the olfactory epithelium normalized to control (log 10, Mean ± SEM, n=4, *p<0.01 (Mann-Whitney test)).

### Damage of the infected olfactory epithelium is correlated with infiltration of innate immune cells

Since apoptosis does not significantly occur in the OE during the initial phase of infection, the desquamation of the infected OE may be related to immune cell infiltration (Bryche et al., 2020; Urata et al., 2021). So far, the immune cells in the nasal cavity have been poorly characterized. Neutrophils and macrophages are known for their importance in clearing infected tissue (*41*) but only Iba1^+^ cells are well characterized in the OE (*21*). Iba1^+^ cells are described as microglia/macrophages but CD68 is more classically used as a marker of monocytes and macrophages (*42*). Concerning neutrophils, the presence of neutrophil cytosol factor 2 (ncf2; (*43*)) and myeloperoxydase (MPO) have been used successfully to characterize these cells in hamsters (*44*).

We first evaluated at 1 dpi and 2 dpi by qPCR the expression of Iba1, CD68 and Ncf2 along with classical inflammatory markers (TNFα and IL6) and the presence of the virus (**Supp. Fig 1**). At 1 dpi, SARS Nucleocapsid protein (SARS N) was already abundantly expressed in the OE at a similar level as at 2 dpi, and TNFα and IL6 transcripts increased gradually (Mann-Whitney, *p*<*0.05*). Iba1 and CD68 expression related to macrophage presence in the OE did not rise significantly at 1 dpi compared to control (Mann-Whitney, *p*=*0.164* and *0.128* respectively) but did at 2 dpi (Mann-Whitney, *p*<*0.05*). Concerning neutrophils, ncf2 expression was strongly enhanced at 1 dpi and was still increasing at 2 dpi (Mann-Whitney, *p*<0.05). These results suggest that neutrophils are already recruited at 1 dpi and that their recruitment continues at 2 dpi along with the arrival of Iba1^+^ and CD68^+^ cells.

We next focused on immunostaining to characterize the presence of Iba1^+^, CD68^+^ and MPO^+^ cells. In the OE of an uninfected hamster, Iba1^+^ cells were already present and mainly localized in the lamina propria while CD68 signal was absent (**Fig. 2A**) indicating that Iba1^+^ cells do not express the classical CD68 marker of macrophages. This was confirmed in the infected areas of the OE where we observed a very different presence of Iba1^+^ and CD68^+^ cells. Iba1^+^ cells were massively present as soon as 1 dpi in the damaged parts of the infected OE as well as in the desquamated cells in the lumen of the nasal cavity. CD68^+^ cells were less abundant in the damaged part of the OE and mainly present in the desquamated cells filling the lumen of the nasal cavity (**Fig. 2B**). A double staining against Iba1 and CD68 of the desquamated cells in the lumen of the nasal cavity did not reveal any overlap of the two markers (**Supp. Fig. 2)**, showing that Iba1^+^ cells do not express CD68 once they are located among the desquamated cells. Similar to CD68, MPO signal was absent in uninfected OE and appears partly in the damaged OE and mainly in the lumen of the nasal cavity along with desquamated cells. Overall, these results show that Iba1^+^ microglia are much more abundant in the infected OE than CD68^+^ macrophages and MPO^+^ neutrophils cells, both being mainly present in the desquamated cells filling the lumen of the SARS-CoV-2 infected nasal cavity.

**Figure 2:**
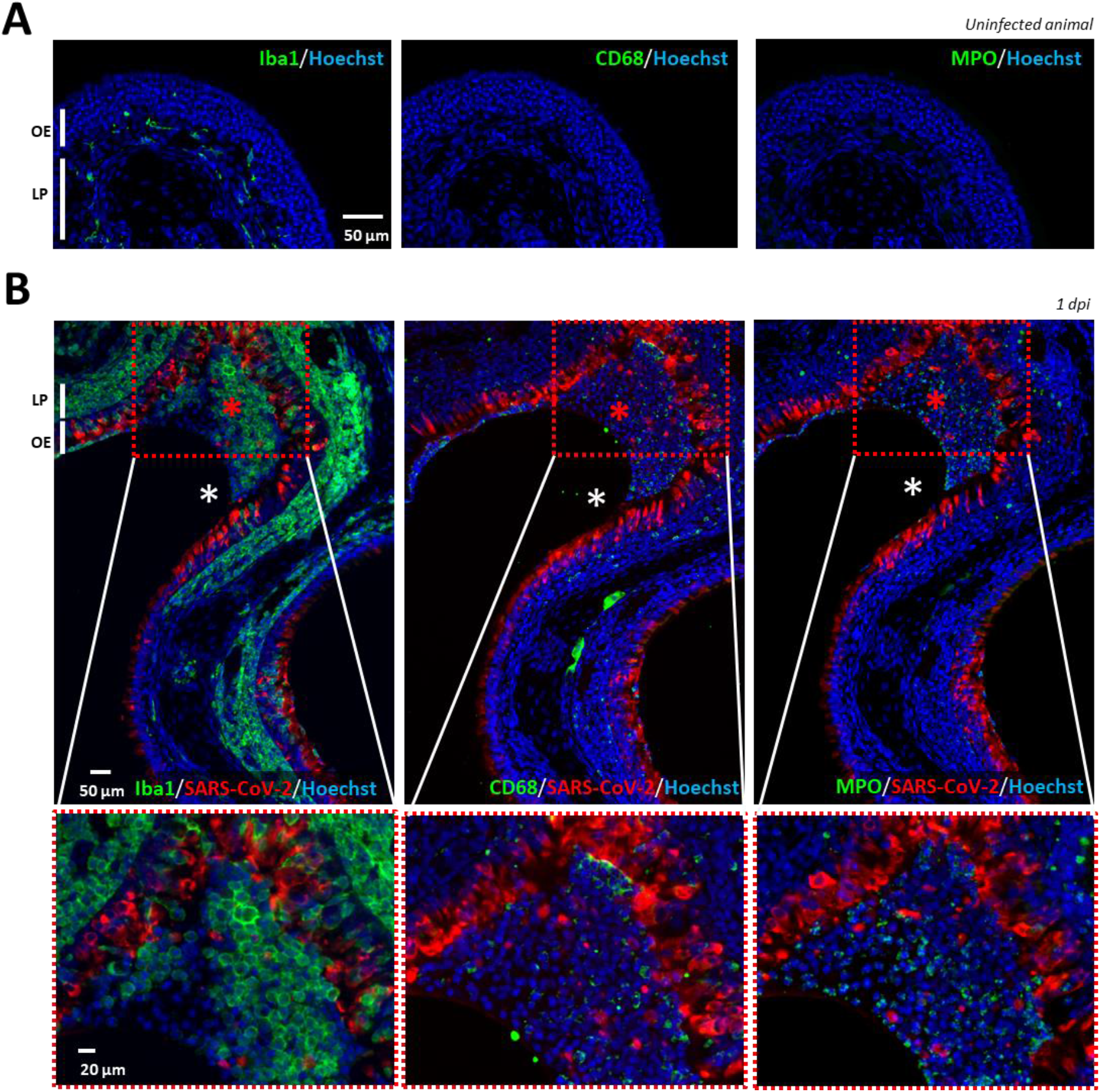
Iba1^+^ (microglia), CD68^+^ (macrophages) and MPO^+^ (neutrophils) cells presence in the olfactory epithelium before and during SARS-CoV-2 infection. Immunostaining on successive slides of the olfactory epithelium from a non-infected (**A**) or 1 dpi hamster (**B**). Only Iba1^+^ cells are present in the uninfected olfactory epithelium (OE) and in the lamina propria (LP). In the infected epithelium, Iba1^+^ cells are massively present in the OE while CD68^+^ and MPO cells are mostly present in the desquamated cells (red asterisk) in the lumen of the nasal cavity (white asterisk).

If these innate immunity cells are involved in the desquamation of the OE, we should always observe their presence in the damaged infected area of the OE. To investigate their infiltration in the OE and its correlation with damage, we focused on 3 zones similarly as for apoptosis quantification: ^2/^uninfected, infected ^2/^without or ^3/^with damage at 1 and 2 dpi. The infiltration level of Iba1^+^ cells in the OE was increased in the damaged infected zone but not in the undamaged one (**Fig. 3**). This difference was statistically significant at 2 dpi (Mann-Whitney, *p*= *0.0286*) and nearly significant at 1 dpi (*p*= *0.0525*). The infiltration of these cells was similarly increased in the lamina propria underneath the previous OE zones with a significant difference at 2 dpi (*p*= *0.0286*). We observed a significant correlation between the damage of the OE and their presence in both the OE and the underlying lamina propria (Spearman test, *p* = *0.0098* and *0.0006* respectively).

**Figure 3:**
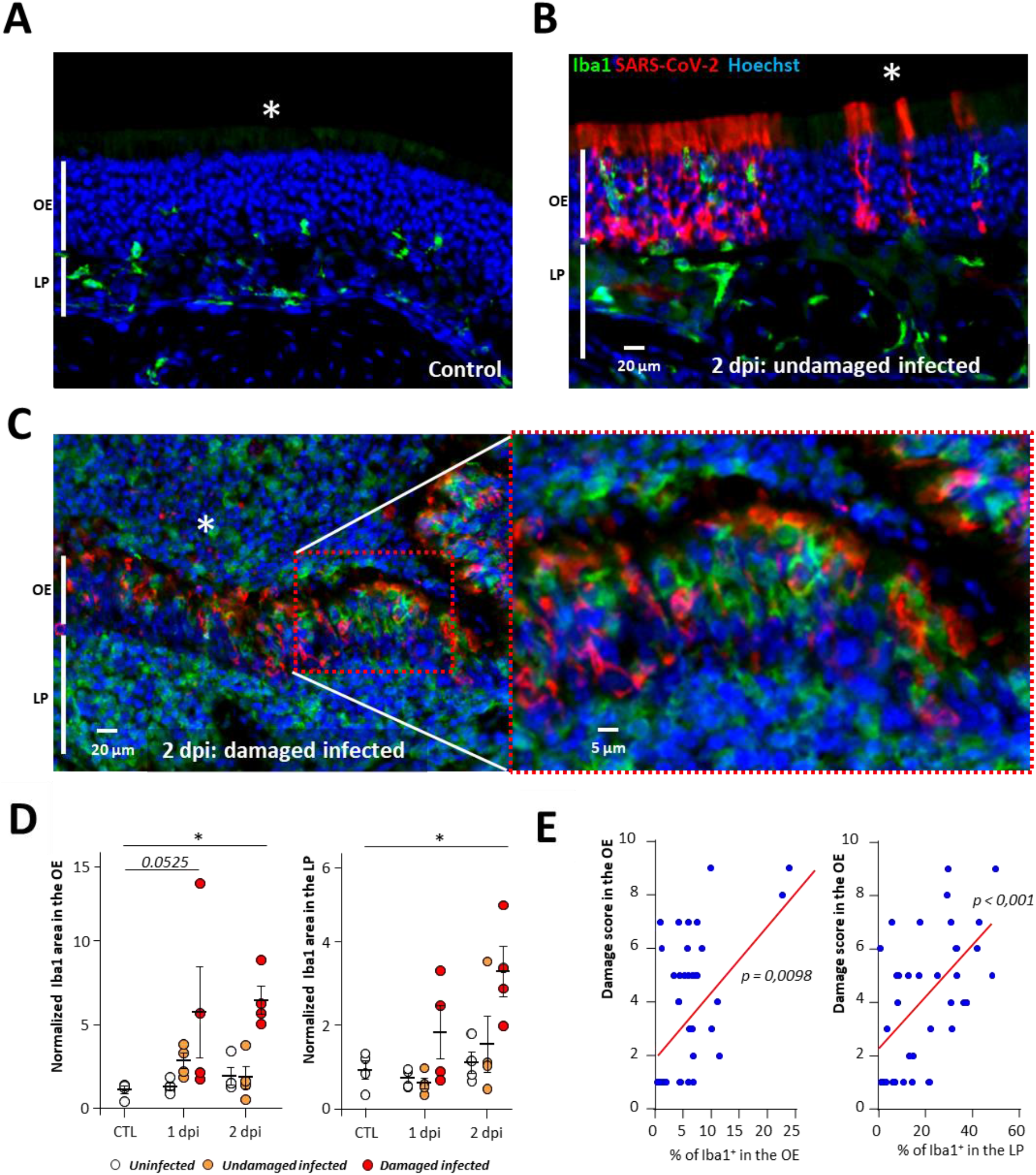
Iba1^+^ cell infiltration increases with the damage in the OE. Representative images of uninfected (**A**), infected but undamaged (**B**) and infected and damaged (**C**) area of the olfactory epithelium (OE) at 2 days post infection (dpi). The lumen of the nasal cavity is indicated by a white asterisk. (**D**) Iba1^+^ signal in the olfactory epithelium (OE, left) and lamina propria (LP, right) in either control animals (CTL) or at 1 or 2 dpi (Mean normalized to control ± SEM, n=4, *p<0.01 (Mann-Whitney test)). (**E**) Correlation between score damage of the olfactory epithelium and the percentage of Iba1^+^ signal in the olfactory epithelium (left panel) and the lamina propria (right panel). Spearman test p value.

We similarly examined whether the presence of CD68^+^ macrophages and MPO^+^ neutrophils in the OE was associated with the damage of the OE after SARS-CoV-2 infection. Both CD68 and MPO signals were increased in the damaged infected zone but not in the undamaged one (**Fig. 4**). This difference was statistically significant in the infected damaged zones at 1 and 2 dpi for both markers compared to control and infected undamaged zones of the OE and lamina propria (Mann-Whitney, p< 0.05). We observed a significant correlation between the damage of the OE and their presence the presence of both CD68^+^ and MPO^+^ cells the OE and the lamina propria (Spearman test, p <0.001).

**Figure 4:**
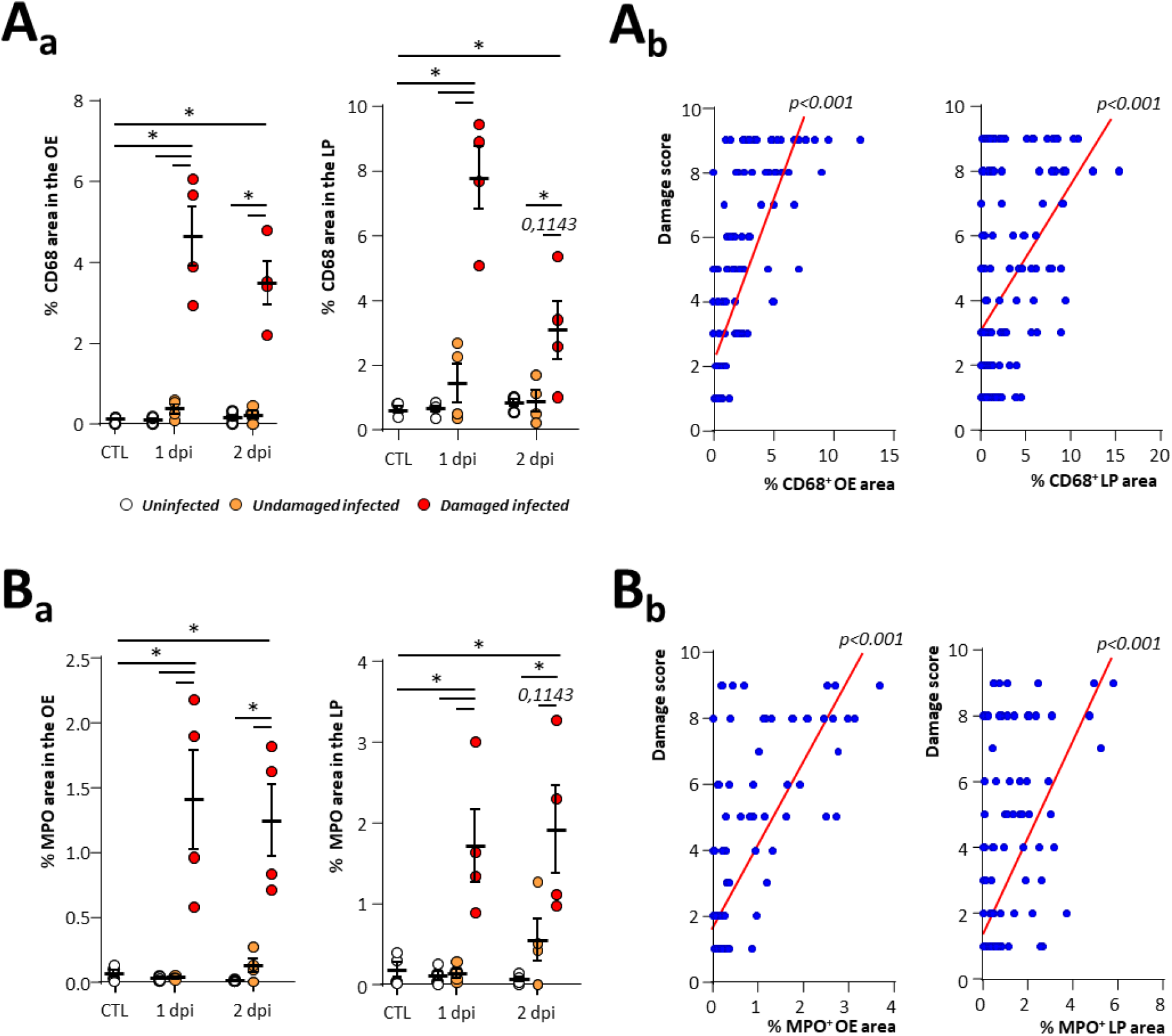
CD68^+^ macrophage and MPO^+^ neutrophil cells are associated with damage of the olfactory epithelium during SARS-CoV-2 infection. CD68^+^ (**A_a_**) and MPO^+^ (**B_b_**) signal in the olfactory epithelium (OE, left) and lamina propria (LP, right) in either control animals (CTL) or at 1 or 2-days post infection (dpi) (Mean normalized to control ± SEM, n=4, *p<0.05 (Mann-Whitney test)). Correlation between score damage and percentage of CD68^+^ (**A_a_**) and MPO ^+^ (**B_b_**) signal in the olfactory epithelium (left panel) and the lamina propria (right panel). Spearman test p value.

### Neutropenia reduces damage of the OE related to SARS-CoV-2 infection as well as level of virus infection

Neutrophils are the main actors of damage to the olfactory epithelium during Poly(I:C) induced inflammation (*45*). We therefore evaluated whether a neutropenic treatment based on cyclophosphamide would reduce the damage induced by SARS-CoV-2 infection in the OE. Such treatment causes apoptosis of bone marrow derived cells and has previously been used successfully on hamsters to induce neutropenia (*44*). We first monitored in control animals how the treatment impacts circulating immune cells. Neutrophils population was decreased by ~10-fold, lymphocytes were also decreased by ~3-fold, and monocytes by ~5-fold (**Supp Fig. 3A**). Since cyclophosphamide can impact basal cell proliferation and thus OE structure, we examined its effect in uninfected animals and did not observe any evident damage on the OE structure (**Supp Fig. 3B**). We next examined the impact of this treatment on the expression of genes related to the innate immune system in the nasal turbinates during SARS-CoV-2 infection. At 1 dpi, the expression of Iba1 and CD68 reflecting the population of microglia and monocytes/macrophages respectively was not statistically different between vehicle and cyclophosphamide treated animals but a decrease of Ncf2 expression reflecting a reduced presence of neutrophils almost reached significance (Mann Whitney; *p*=*0.0571*; **Fig. 6A**). We observed a tendency of TNFα and IL6 expression reduction which did not reach significance either (*p*=*0.1143*). Despite the overall tendency of a decrease of innate immune system response at 1 dpi, the level of SARS-CoV-2 infection reflected by N protein expression was decreased and the difference almost reached significance (*p*=*0.0571*). At 2 dpi, the expression of all genes related to innate immune cell presence as well as TNFα were decreased (*p*<*0.05*). We could not make definitive conclusions about IL6 and SARS-CoV-2 N protein expression because it was very variable in cyclophosphamide treated animals. We examined histologically the damage and the level of infection in the OE (**Fig. 5 B, C**). MPO signal was significantly decreased at 1 dpi in the OE of cyclophosphamide treated animals compared to control (*p*<*0.05*) but was too variable at 2 dpi to reach significance (**Fig. 5 D_a_**). The damage in the OE was significantly decreased at 1 and 2 dpi (*p*<*0.01* and *p*<*0.05* respectively) (**Fig. 5 D_b_**). The reduction tendency of the virus presence measured by the N protein expression was confirmed by immunostaining in the OE at 1 dpi only (*p*<*0.05;***Fig. 6 D_c_**). We hypothesize that this reduction could be linked to less infected desquamated cells released into the lumen of the nasal cavity following the damage induced by the neutrophils. We thus quantified the area of desquamated cells in the lumen which was significantly decreased at 1 dpi and almost reached significance at 2 dpi (**Fig. 5 D_d_**; *p*<*0.001* and *p*=*0.0905* respectively). In the desquamated cells of the lumen, the percentage of infected cells was significantly diminished at 1 dpi (**Fig. 5 D_e_**; *p*<*0.001*) but not at 2 dpi when infected desquamated cells were in the lumen of the nasal cavity in some treated animals (**Supp Fig. 3C**). Since the reduction of SARS-CoV-2 replication in the nasal cavity of immunocompromised animal was unexpected, we verified that a dose 3 times higher than the maximum dose of cyclophosphamide potentially present in the hamsters during infection did not limit the virus replication *in vitro* (**Supp Fig. 4A**).

**Figure 5:**
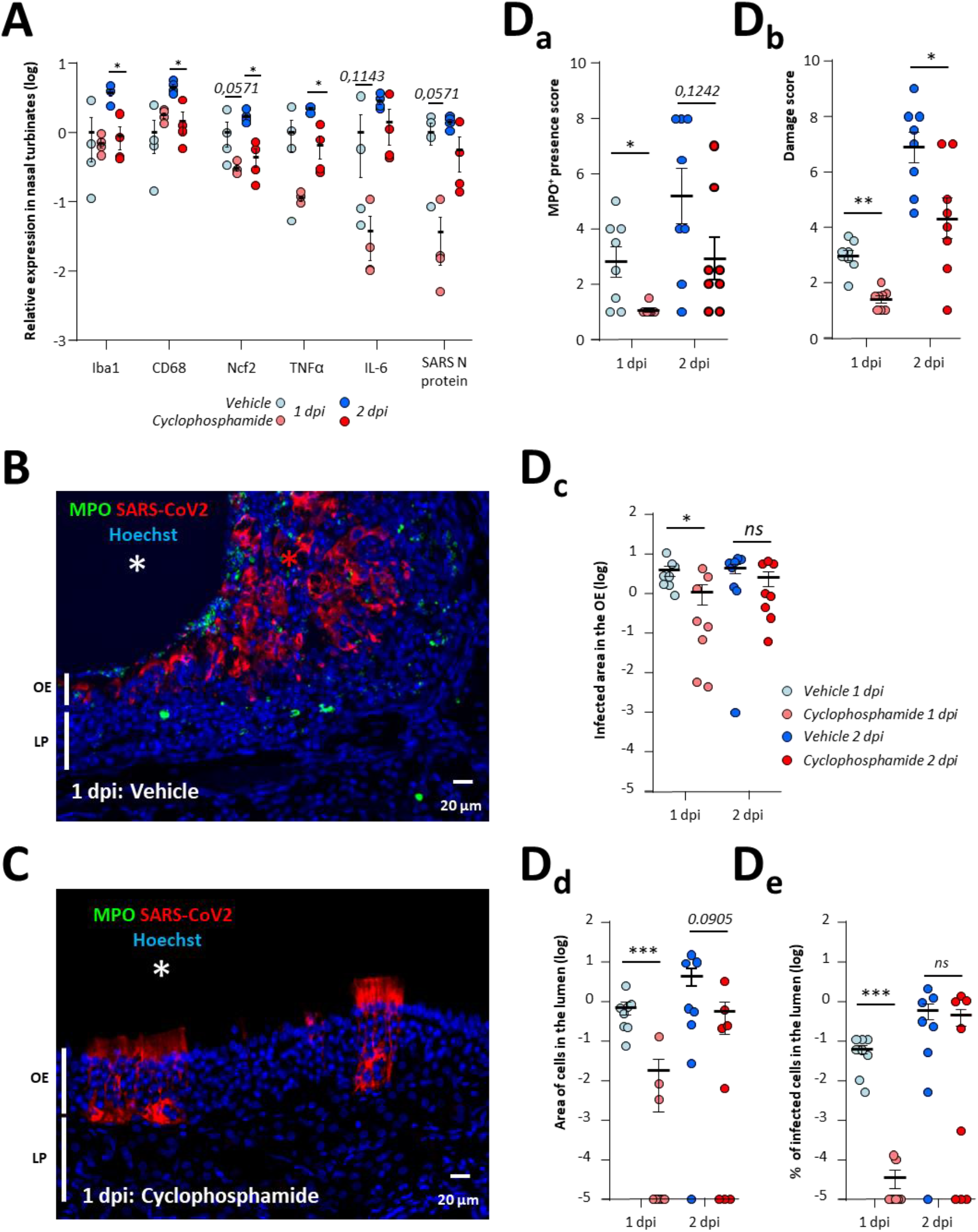
Immunosuppression induced by cyclophosphamide reduces damage of the olfactory epithelium as well as virus replication. (**A**) Expression of innate immune genes in the nasal turbinates with or without cyclophosphamide treatment at 1 and 2-days post infection (dpi). Iba1, CD68 and Ncf2 are related to the presence of microglia/macrophages, monocytes/macrophages and neutrophils respectively; TNFα and IL6 are two cytokines expressed during inflammation; SARS-CoV-2 N expression is related to the SARS-CoV-2 infection. Results represent the mean ± SEM relative to vehicle treated hamsters (n=4, *p<0.05; Mann-Whitney test). Representative images of the infected olfactory epithelium immunostained for MPO (neutrophil marker) and SARS-CoV-2 N protein in (**B**) vehicle and (**C**) cyclophosphamide treated animal (olfactory epithelium (OE), lamina propria (LP)). In the vehicle condition, the lumen (white asterisk) is filled with desquamated cells (red asterisk) containing MPO signal. In the cyclophosphamide condition, MPO signal is absent and the lumen is mostly free of cellular debris. Quantification in the OE of (**D**_1_) MPO^+^ neutrophil presence (**D**_2_) damage score (**D**_3_) SARS-CoV-2 infected area and in the lumen of the nasal cavity of (**D**_4_) desquamated cells area and (**D_5_**) percentage of SARS-CoV-2 infected area in the desquamated cells (Mean normalized to infected animals treated with vehicle ± SEM, n=8 areas of the nasal cavity from 4 different animals, *p<0.05, **p<0.01, ***p<0.001 (Mann-Whitney)).

**Figure 6:**
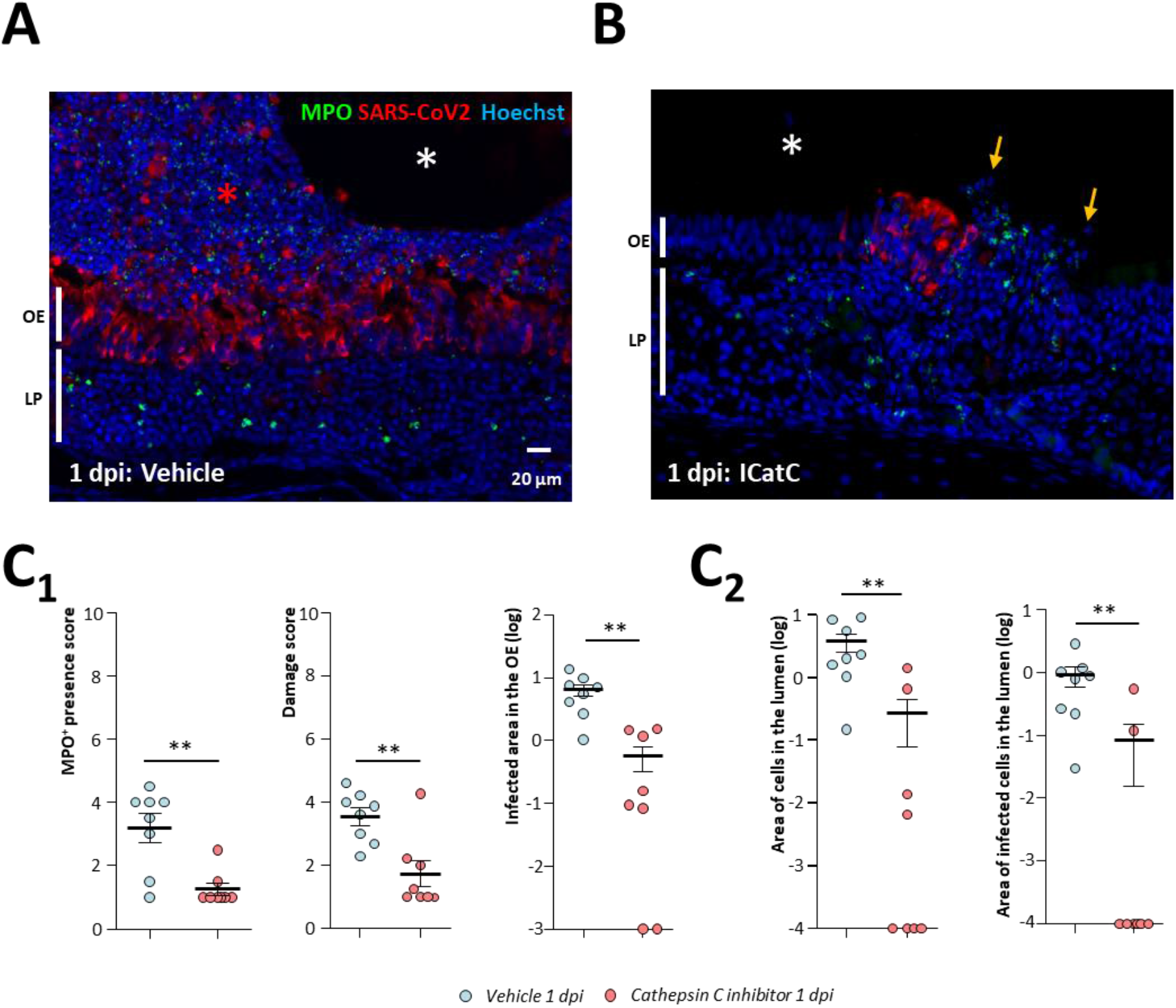
Inhibition of neutrophil proteinases reduces damage of the olfactory epithelium as well as virus spreading in the nasal cavity. Representative images of the infected olfactory epithelium immunostained for MPO (neutrophil marker) and SARS-CoV-2 N protein in (**A**) vehicle and (**B**) cathepsin C inhibitor (Icat_CXPZ-01_) treated animal (olfactory epithelium (OE), lamina propria (LP)). In the vehicle condition, the lumen (white asterisk) is filled with desquamated cells (red asterisk) containing MPO signal. Under cathepsin C inhibition, MPO signal is less abundant and the lumen is mostly free of cellular debris (yellow arrows). Quantification **(C_1_)** in the OE of MPO^+^ neutrophil presence; damage score; SARS-CoV-2 infected area and **(C_2_)** in the lumen of the nasal cavity of desquamated cell area and percentage of SARS-CoV-2 infected area in the desquamated cells (Mean normalized to infected animals treated with vehicle ± SEM, n=8 areas of the nasal cavity from 4 different animals, *p<0.05, **p<0.01, ***p<0.001 (Mann-Whitney test)).

### Inhibition of neutrophil proteinases reduces damage of the OE related to SARS-CoV-2 infection as well as virus spreading in the nasal cavity

In order to confirm our results on cyclophosphamide treatment which affects immune cells other than neutrophils, we treated animals with an inhibitor of cathepsin C (IcatC_XPZ-01_) which is essential for the maturation of elastase-like proteinases of neutrophils. This inhibitor has been used successfully to almost completely eliminate the elastase-like activity of neutrophils *in vivo* (*38*). We chose to focus on the histological impact of IcatC_XPZ-01_ treatment at 1 dpi as it gave the most significant results during cyclophosphamide treatment. The inhibition of elastase-like proteinases of neutrophils gave similar results as cyclophosphamide treatment even though we sometimes observed locally restricted neutrophil infiltration in the OE (**Fig. 6 A, B**). Globally the MPO^+^ neutrophil presence in the nasal turbinates and the damage in the infected area of the OE were significantly reduced compared to vehicle treated animals (**Fig. 6C_1_**; Mann-Whitney; *p*<*0.01*). We also observed significantly less desquamated cells in the lumen of the nasal cavity which were also less infected (**Fig. 6C_2_**; Mann-Whitney; *p*<*0.01*). Since we observed again that the inhibition of neutrophil action limited SARS-CoV-2 replication, we verified that a dose 3 times higher than the maximum dose of IcatC_XPZ-01_ potentially present in the hamsters during infection did not impair the virus replication *in vitro* (**Supp Fig. 4B**).

## Discussion

The anosmia induced by SARS-CoV-2 infection is now clearly linked to the infection of the olfactory epithelium with a main tropism for sustentacular cells (*46*, *47*). We and others have observed that following this infection, the OE undergoes massive damage leading to cell desquamation and cellular debris filling the lumen of the nasal cavity (*13*, *19*, *20*), but the mechanism of this destruction is less clear.

Several studies reported an increase of apoptosis especially in olfactory sensory neurons (*18*–*20*) and assumed that it led to the destruction of the OE. Here, we first examined the apoptosis level based on cleaved caspase 3 level in the OE of uninfected animals and infected areas of the OE, either intact or damaged. If apoptosis initiates the desquamation process then it should increase in the damaged areas of the OE. However, we observed a similar level of apoptosis level in all these areas which was consistent with the basal level of apoptosis that we previously measured in adult mice and rats OE (*48*, *49*). While the apoptosis in the infected damaged OE was low, we observed an increased level of apoptosis in cells present in the lumen of the nasal cavity. The discrepancy with previous studies may be due to different models as some were performed using transgenic mice expressing hACE2 but it should be noted that these studies did not perform any quantification or misinterpreted some apoptosis staining in the lumen of the nasal cavity as can be observed with TUNEL staining (*20*). The induction of apoptosis by loss of cell contact is well described (*50*), a phenomenon known as anoikis (*51*). It indicates that desquamated cells are sufficiently preserved to be able to enter apoptosis after the desquamation process is initiated following OE SARS-CoV-2 infection.

Since we previously observed that the infected area of the OE is infiltrated by immune cells (*13*), we next explored whether innate immune cells are involved in this process, especially macrophages and neutrophils known to be involved in damage of epithelial cells during acute inflammation (*28*, *29*). If so, we should systematically observe their presence in the damaged area of the OE. We first characterized the presence of these cells in the infected OE. We observed that CD68, a classical marker of macrophages (*42*) was expressed in a different population than Iba1^+^ cells previously described as a microglia/macrophages cellular population (*24*). Therefore, in the following, we will refer to Iba1^+^ cells as microglia, and CD68^+^ cells as macrophages to avoid confusion between these two cells types. Immunostaining revealed that at 1 dpi, some parts of the OE in the most rostral part of the nasal cavity were already significantly damaged. We observed that in these tissues, microglia cells were mainly recruited while macrophages and neutrophils appeared in the zones desquamating toward the lumen of the nasal cavity. We can thus hypothesize that microglia are first infiltrating the SARS-CoV-2 infected OE. This is not consistent with our qPCR results indicating that neutrophils are recruited more abundantly at 1 dpi than microglia and macrophages. However, we observed that contrary to macrophages and neutrophils, microglia cells were already present in the lamina propria of uninfected zones. At the beginning of infection, the increase of microglia cells could simply arise from infiltration of adjacent cells in the nasal turbinates, while macrophages and neutrophils may migrate from the blood following chemotaxis. Microglia are recruited around infected cells of the central nervous system within hours (Fekete et al., 2018) which is consistent with our observation in the OE. Further studies are required to decipher the origin of these 3 cells population during the early events of SARS-CoV-2 infection in the infected nasal turbinates.

Neutrophils are known to induce epithelial damage and an elegant study demonstrated their major role during Poly I:C (an artificial double stranded RNA agonist of TLR3 receptor) nasal instillation leading to damage of the OE (*45*). In order to evaluate the importance of the neutrophils in the damage induced by SARS-CoV-2, we first induced neutropenia based on previous cyclophosphamide treatment successfully used in hamsters (*44*). We confirmed that such treatment mainly impacts neutrophils but also reduced to a lesser degree other leucocyte populations. As expected such treatment reduced neutrophil infiltration in infected areas of the OE and we observed that damage of the infected OE was significantly reduced as well. In order to confirm the role of neutrophils in the damage of the OE following SARS-CoV-2 infection, we treated hamsters with an inhibitor of cathepsin C (IcatC_XPZ-01_) strongly reducing the neutrophil elastase-like proteinases activity. We observed that similar to cyclophosphamide treatment, the damage of the OE was greatly reduced in this context. Surprisingly, the global infiltration of neutrophils was reduced as well, even though we observed that some neutrophils were still present in the most infected area of the OE. Since neutrophils are mainly present among the desquamated cells present in the lumen of the nasal cavity, the reduction in neutrophil infiltration during IcatC_XPZ-01_ treatment may be linked to a decrease of the OE damage as elastase-like proteinase action increases inflammation (*52*). It suggests that the damage of the OE initiated by neutrophils may participate in the increasing infiltration of neutrophils leading *in fine* to massive damage of the infected OE areas. Overall, these results show that neutrophils have a major causative role in the destruction of the OE following SARS-CoV-2 infection by releasing elastase-like proteinases.

We observed that at 1 dpi, the level of SARS-CoV-2 infection was reduced in cyclophosphamide and IcatC_XPZ-01_ treated hamsters. Such a result was unexpected as neutrophils should effectively destroy infected cells and thus their action should reduce infection progression. Since we observed that these treatments did not impair SARS-CoV-2 replication *in vitro*, we hypothesize that neutrophils may have a counterproductive effect by releasing infected cells into the lumen of the nasal cavity. These infected cells could allow the virus to spread more easily in the nasal cavity due to impairment of mucociliary clearance that has been recently shown to be reduced during SARS-CoV-2 infection (*53*). Such a hypothesis is consistent with our results showing that cyclophosphamide and IcatC_XPZ-01_ treatment significantly reduced the amount of infected desquamated cells filling the lumen of the nasal cavity. Such a phenomenon has been observed recently *in vitro* in respiratory epithelial cell culture where infection is enhanced in the presence of neutrophils. In their preliminary study, SARS-CoV-2 alone did not significantly increase cytokine production but the neutrophil presence did, showing a major role of the neutrophils in the epithelial response to SARS-CoV-2 infection (*54*). At 2 dpi, the impact of neutropenia on SARS-CoV-2 infection in the nasal cavity was statistically significant through the reduction of OE damage. The neutropenia induced by the cyclophosmamide treatment was only partial, we can thus hypothesize that the remaining neutrophils can be more effectively recruited when infection progresses and thus we could no longer observe a statistically significant decrease of neutrophil infiltration and SARS-CoV-2 infection in the OE.

Overall, our results show that the SARS-CoV-2 infection does not directly induce the massive damage of the OE but that neutrophils play a major role by releasing elastase-like proteinase in the infected OE. This probably leads to the destabilisation of the OE structures and the expulsion of cells including non-infected neurons into the lumen of the nasal cavity where they would undergo apoptosis. In the early phase of the infection, such expulsion of infected cells could enhance the virus infection in the nasal cavity (**Fig. 7**). We observed damaged areas of the OE as soon as 1 dpi indicating that the innate immune system is extremely efficient in detecting SARS-CoV-2 infected cells to destroy them. The signal triggering this very fast action remains to be explored. The host’s immune defence system that may be present to prevent pathogen invasion from the nose to the brain seems beneficial for SARS-CoV-2 to achieve a much more extensive infection of the OE than any previous virus, resulting in unprecedented olfactory dysfunction in the COVID-19 pandemic.

**Figure 7:**
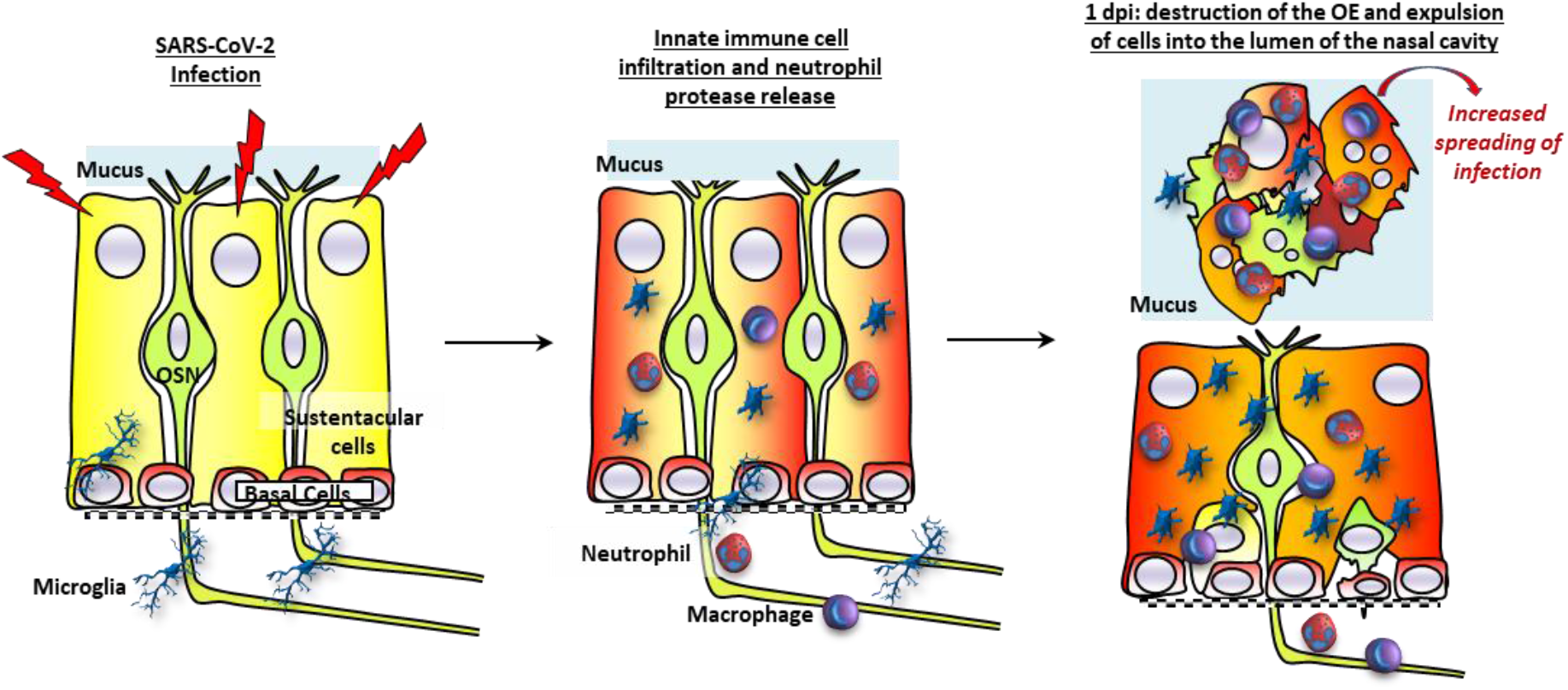
Model of innate immune cell signalling leading to olfactory epithelium desquamation. The olfactory epithelium is mainly composed of olfactory sensory neurons (OSN) surrounded by supporting cells (sustentacular cells) and basal cells able to regenerate all cell types of the epithelium. During the infection of sustentacular cells (turning red), microglia become activated and infiltrate the olfactory epithelium followed by neutrophils and macrophages. Neutrophils release elastase-like proteinase leading to destabilization of the epithelium structures and the expulsion of cells including non-infected neurons into the lumen of the nasal cavity. These desquamated cells – after losing cell contact – undergo apoptosis with a nuclear fragmentation. The release of infected cells may contribute to an increased spreading of the virus in the nasal cavity.

## Abbreviations

OE: (olfactory epithelium)
OSN: (olfactory sensory neuron)
SCs: (sustentacular cells)
MPO: (myeloperoxidase), neutrophil, proteases

## Supplementary materials

**Supplementary Table 1**: Sequences of primers

**Supplementary Figure 1:** Expression of different genes in the nasal turbinates during SARS-CoV-2 infection

**Supplementary Figure 2**: Double staining against Iba1 and CD68 markers in the desquamated cells filling the lumen of the nasal cavity

**Supplementary Figure 3**: Blood numeration, impact on OE structure and representative images of the nasal cavity at 2 dpi during cyclophosphamide treatment.

**Supplementary Figure 4**: Impact of cyclophosphamide and cathepsin C inhibitor on virus replication *in vitro*.

## Acknowledgment

We would like to thank all VIM members for their helpful discussion, Christopher von Bartheld and Birte Nielsen for improvement of the manuscript, Dr Pierre Deshuillers from the BioPole plateform of the National Veterinary School of Alfort for hamsters’ blood count, all the people from the PRBM plateform of ENVA who helped us in the BSL3 animal facility. Bertrand Bryche, Georges Saade, Mustapha Si-Tahar and Déborah Diakite for helpful discussion and technical help.

## Authors contribution

Conceptualization NM and CB Investigation: CB, ASA, OAG, BDC, RD, BK, BK, SLP, NM Formal analysis: CB and NM Writing: NM and CB with input from all authors.

