## Supplemental data for "Neutrophils initiate the destruction of the olfactory epithelium during SARS-CoV-2 infection in hamsters"

| Gene | Primer 5’>3’ | Primer 3’>5’ |
|---|---|---|
| G3PDH | GACATCAAGAAGGTGGTGAAGCA | CATCAAAGGTGGAAGAGTGGGA |
| SARS 2 N | GGGGAACTTCTCCTGCTAGAAT | CAGACATTTTGCTCTCAAGCTG |
| Iba1 | TGGATGAGATCAACAAGCAATTC | AAGGCTTCCAGTTTGGAGGG |
| CD68 | CACTTGGGGCCATGTTTCTC | CTCGGGTAATGCAGAAGGCA |
| Ncf2 | ATGTTCAATGGACAGAAGGGGC | TGGGATCTTTCTGGGGCACT |
| TNF-α | AACTCCAGCCGGTGCCTAT | GTTCAGCAGGCAGAAGAGGATT |
| IL-6 | AGACAAAGCCAGAGTCATT | TCGGTATGCTAAGGCACAG |

**Supplementary Table 1:** Sequences of primers

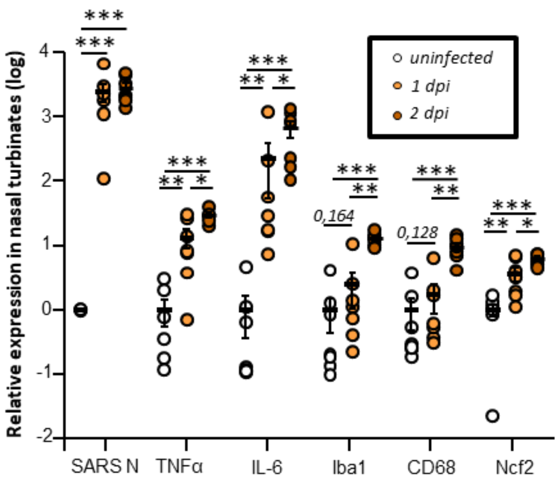


***Supplementary Figure 1: Expression of different genes in the nasal turbinates during SARS-CoV-2 infection.*** *SARS-CoV-2 N expression is related to the SARS-CoV-2 infection ; TNFα and IL6 are two cytokines expressed during inflammation ; Iba1, CD68 and ncf2 are related to the presence of microglia/macrophages, activated monocytes-derived macrophages and neutrophils respectively. Results represent the mean ± SEM relative to uninfected hamsters (n=7, *p<0.05, **p<0.01, ***p<0.001; Mann-Whitney test).*

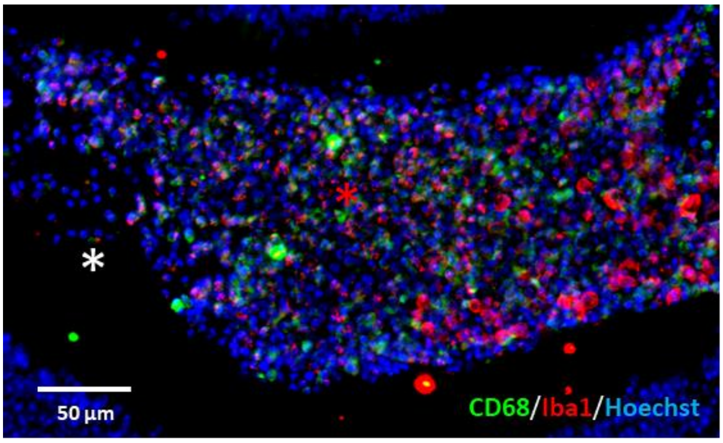


**Supplementary Figure 2:** Double staining against Iba1 and CD68 markers in the desquamated cells (red asterisk) filling the lumen of the nasal cavity (white asterisk) of a SARS-CoV-2 infected hamster at 2 dpi. Part of the desquamated cells are Iba1$^{+}$ and CD68$^{+}$ with no co-staining.

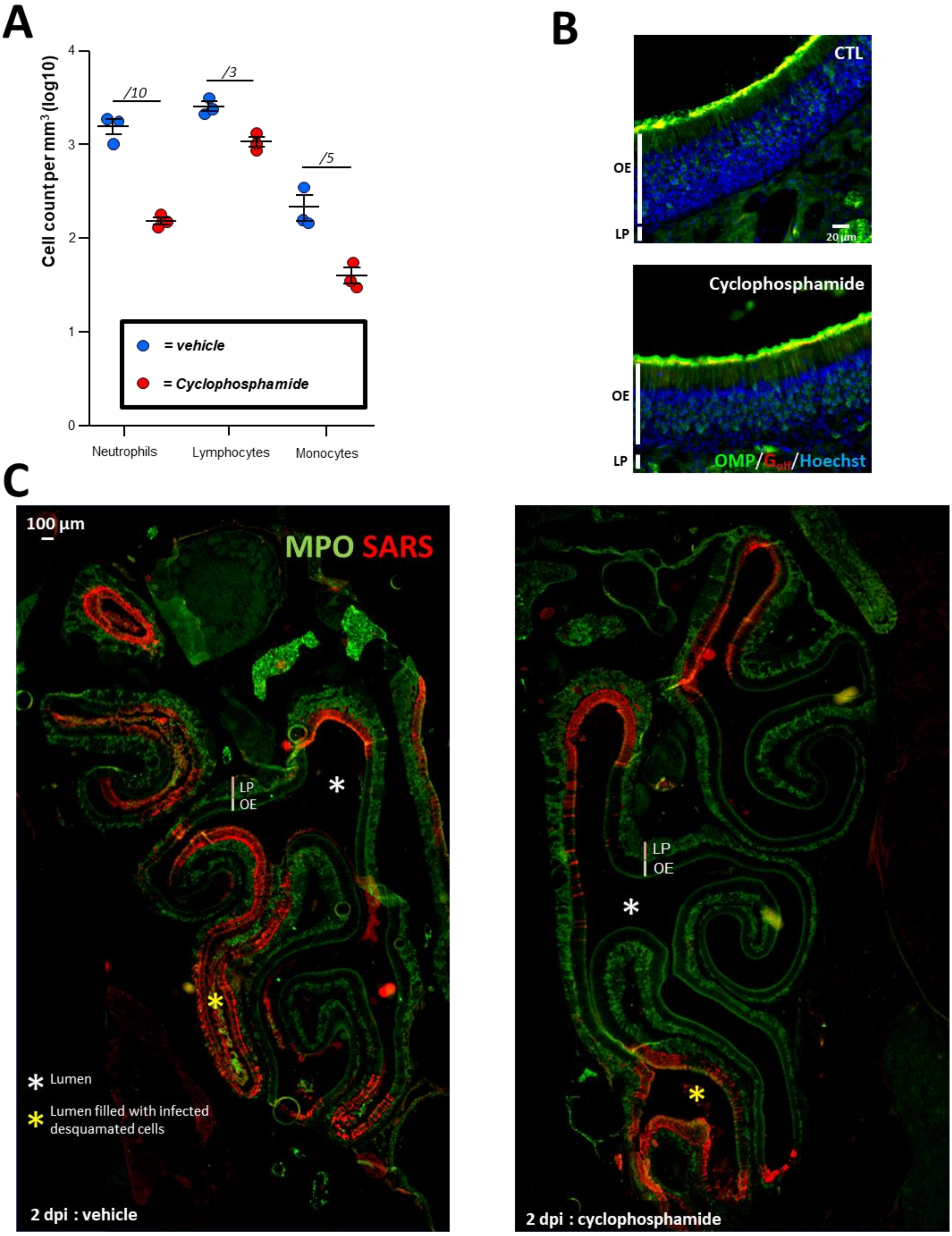


**Supplementary Figure 3:** (**A**) Blood numeration of neutrophils, lymphocytes and monocytes, the number above the data points represents the difference between the mean of the control *vs* treated (Mean ± SEM, n=3) (**B**) Representatives images of the olfactory epithelium immunostained against OMP (Olfactory Marker Protein staining mature olfactory neurons) and olfactory specific G protein ($G_{olf}$ present in cilia of olfactory neurons) in control (CTL) or cyclophosphamide treated uninfected animals. (**C**) Global view of the nasal cavity at 2 dpi showing infection level in the OE immunostained against SARS-CoV-2 N protein (red), in cyclophosphamide treated animals, the level of infection is reduced but there are some desquamated infected cells (yellow asterisk) present into the lumen of the nasal cavity (white asterisk).

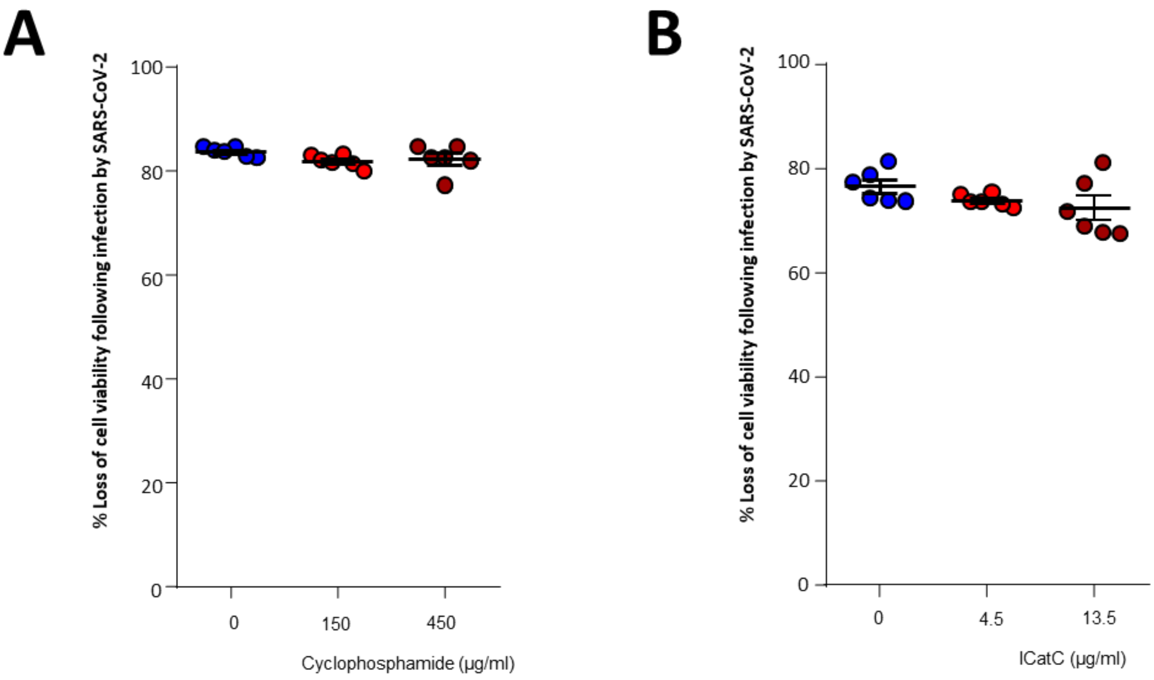


**Supplementary Figure 4:** Impact of (**A**) cyclophosphamide and (**B**) Cathepsin C inhibitor (ICatC) on Vero E6 cell death following SARS-CoV-2 infection (n=6, mean ± SEM).
